# KIF24 controls the clustering of supernumerary centrosomes in pancreatic ductal adenocarcinoma cells

**DOI:** 10.1101/2022.03.16.484562

**Authors:** Yu Mashima, Hayato Nohira, Hiroki Sugihara, Brian David Dynlacht, Tetsuo Kobayashi, Hiroshi Itoh

## Abstract

Clustering of supernumerary centrosomes, potentially leading to cell survival and chromosomal instability, is frequently observed in cancers. However, the molecular mechanisms by which centrosome clustering is controlled in cancer cells remain largely unknown. A centrosomal kinesin, KIF24, was previously shown to restrain the assembly of primary cilia in mammalian cells. Here, we revealed that KIF24 depletion suppresses multipolar spindle formation by clustering centrosomes in pancreatic ductal adenocarcinoma (PDAC) cells harboring supernumerary centrosomes. KIF24 depletion also induced hyperproliferation and improved the mitotic progression in PDAC cells. On the other hand, disruption of primary cilia failed to affect the proliferation and spindle formation in KIF24-depleted cells. These results represent a novel role of KIF24 in suppressing centrosome clustering independent of primary ciliation in centrosome-amplified PDAC cells.

## Introduction

The centrosome comprises centrioles and a pericentriolar matrix (PCM). The two cylinder-like centrioles in G1 phase are duplicated through S–G2 phase and the two-paired centrioles ensure bipolar spindle formation during mitosis. As new centrioles are duplicated from the older centriole, the differentially-aged centrioles in G1 phase are termed the older mother and the younger daughter centrioles. The mother centriole, characterized by the distal and sub-distal appendages, nucleates the primary cilium during interphase, generally in G0 phase. This single hair-like organelle is ubiquitously expressed in mammalian cells containing specific receptors and channels to receive multiple signaling molecules. Given that both the spindle and primary cilium share the centriole for their assembly, they are thought to be incompatible with each other in normal somatic cells (Kobayashi & Dynlacht, 2011; Sánchez & Dynlacht, 2016).

Contrary to normal cells, the numbers of centrosomes and primary cilia are anomalous in many cancer cells. The primary cilia decrease or disappear in various cancers, which probably cause aberrant signal transduction and cell cycle progression (Eguether & Hahne, 2018; Liu *et al*, 2018). Supernumerary centrosomes are also observed in cancers that potentially nucleate the multipolar spindles during mitosis, thereby leading to cell death. Despite this, the cancer cells can often avoid the detrimental multipolar spindles by forming pseudo bi-polar spindles in which the multiple centrosomes are clustered into one spindle to allow bi-polar separation, a process termed centrosome clustering (Funk *et al*, 2016; Levine & Holland, 2018; Milunović-Jevtić *et al*, 2016). Pseudo bi-polar mitosis not only allows completion of cell division and survival of the daughter cells, but also occasionally induces certain tumor cell hallmarks, such as chromosome segregation errors, aneuploidies, and invasion (Ganem *et al*, 2009).

Pancreatic ductal adenocarcinoma (PDAC) accounts for over 90% of pancreatic cancers, with a 5-year survival rate of less than 10% (Adamska *et al*, 2017). Similar to other cancers, primary cilia are decreased in PDAC lesions and cell lines (Schimmack *et al*, 2016; Seeley *et al*, 2009). Most PDAC samples harbor actively mutated oncogenic KRAS, and KRAS signaling is known to inhibit primary ciliogenesis in PDAC (Seeley *et al*., 2009). We have previously revealed that KRAS and a histone deacetylase HDAC2 suppress primary ciliation by regulating Aurora A kinase (AurA) expression in PDAC cells (Kobayashi & Itoh, 2017; Kobayashi *et al*, 2017). Further, the suppression of ciliogenesis has been reported to promote the transformation of normal pancreatic ductal cells into cancer cells (Deng *et al*, 2018). Centrosomal amplification and clustering also occur in PDAC (Ansari *et al*, 2018; Mittal *et al*, 2015; Sato *et al*, 1999; Zhu *et al*, 2005); however, the molecular mechanisms underlying the centrosome clustering remain largely unclear.

Kinesin family member 24 (KIF24) belongs to the Kinesin-13 subfamily with a kinesin domain in its middle region (Miki *et al*, 2005). KIF24 localizes to centrioles and interacts with two centriolar proteins, CP110 and CEP97, which prevent the assembly of the primary cilia in cycling mammalian cells (Kobayashi *et al*, 2011). KIF24 likewise suppresses cilia formation through the dual roles in which KIF24 recruits CP110 to the mother centriole and antagonizes the extension of the ciliary axoneme with its microtubule (MT)-depolymerizing activity in non-transformed cells (Kobayashi *et al*., 2011). NEK2 phosphorylates KIF24, which enhances MT-depolymerizing activity (Kim *et al*, 2015). MPP9 also forms a complex with CP110-CEP97-KIF24 to suppress ciliogenesis (Huang *et al*, 2018).

In this study, we generated KIF24-depleted PDAC cells which assemble increased primary cilia to assess whether forced expression of these organelles restrained the proliferation of PDAC cells. However, KIF24-depleted cells exhibited abnormal, enhanced proliferation compared to control cells. KIF24 depletion was found to suppress the formation of multipolar spindles by clustering excess centrosomes and to improve mitotic progression in PDAC cells. On the other hand, the elimination of primary cilia in KIF24-depleted cells failed to affect both the proliferation as well as centrosome clustering. Moreover, KIF24 depletion specifically blocked the formation of multipolar spindles and induced hyper-proliferation in PDAC cells harboring supernumerary centrosomes. These results represent a novel function of KIF24 in the centrosome clustering, irrespective of primary ciliation in centrosome-amplified PDAC cells.

## Results

### KIF24 depletion restores primary cilia in Panc1 cells

To examine whether KIF24 contributes to the suppression of primary ciliogenesis in PDAC cells, we utilized Panc1 cells, which were originally isolated from a PDAC patient and assemble primary cilia with low frequency (Lieber *et al*, 1975; Nielsen *et al*, 2008). Since the siRNA-mediated acute knockdown of KIF24 induced the formation of primary cilia in Panc1 cells (data not shown), KIF24-mutated Panc1 cells were subsequently generated using the CRISPR/Cas9 method. The sequencing analysis indicated four distinct mutations in Kif24-mutated cells (named Kif24-3 cells), leading to premature stop codons in three alleles (allele A–C) but an amino acid deletion in one allele (allele D) (Figure S1A), suggesting that KIF24 was not completely knocked out in Kif24-3 Panc1 cells. A rescue clone was further established in which ectopic KIF24 was stably expressed in Kif24-3 cells (Kif24-3_KIF24), and control clones harboring the empty vector (EV) (Panc1_EV and Kif24-3_EV) were also generated. Western blotting analysis showed KIF24 expression in Panc1_EV and Kif24-3_KIF24 cells but a substantial decrease in KIF24 in Kif24-3_EV cells (Figure 1A), indicating that KIF24 is drastically depleted in Kif24-3 cells. The assembly of primary cilia was assessed in these cells by immunofluorescence experiments using an anti-ARL13B antibody that specifically recognizes the ciliary membrane. Kif24-3 cells formed significantly more primary cilia than Panc1_EV and Kif24-3_KIF24 cells in both serum-fed (FBS+) and -deprived (FBS-) media (Figure 1B, C), suggesting that KIF24 suppresses primary ciliogenesis in Panc1 cells. In addition, Panc1 cells stably expressing shRNA targeting KIF24 were generated (Figure 1D, S1B). The percentage of cells with primary cilia was significantly increased in two individual shKIF24-expressed cells compared to shControl cells (Figure 1E). These results collectively indicate that KIF24 negatively controls the assembly of primary cilia in Panc1 cells, as expected from previous reports (Kim *et al*., 2015; Kobayashi *et al*., 2011).

**Figure 1.**
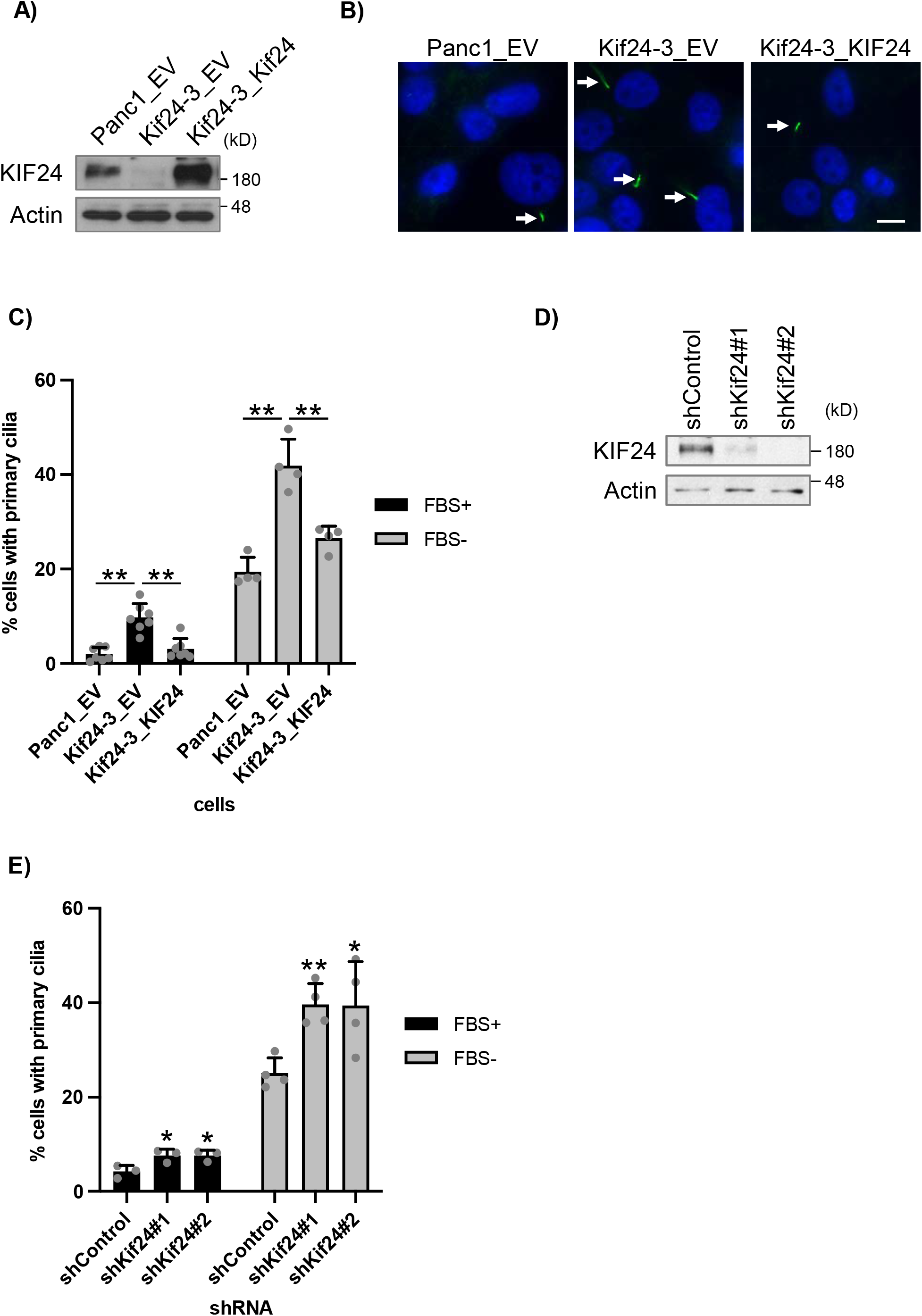
KIF24 depletion restores primary cilia in Panc1 cells. **(A)** The indicated Panc1 cells were cultured in serum-fed medium for 48 hrs. Cell extracts were immunoblotted with an anti-KIF24-2 antibody. β-Actin was used as a loading control. **(B, C)** The indicated Panc1 cells were cultured in serum-fed (FBS+) or serum-starved medium (FBS-) for 48 h and immunostained with an anti-Arl13b antibody (green). (B) Representative images of cells in FBS-cultivation. Arrows indicate primary cilia. DNA was stained with Hoechst (blue). Scale bar, 10 µm. (C) The percentage of ciliated cells was determined. The average of four to seven independent experiments is shown; >250 cells were scored each time. **(D)** The indicated Panc1 cells were cultured and immunoblotted as described in Figure 1A. **(E)** The indicated Panc1 cells were cultured and immunostained as described in Figure 1B. The percentage of ciliated cells was determined. The average of three to four independent experiments is shown; >250 cells were scored each time. **(C, E)** All data are shown as mean ± SD. two-tailed Student’s *t*-test. **, *p* < 0.01; *, *p* < 0.05.

### Loss of KIF24 enhances the proliferation of Panc1 cells *in vitro* and *in vivo*

The proliferation of KIF24-depleted cells was evaluated *in vitro*. As primary cilia appear to inhibit cell division, KIF24-depleted cells were predicted to exhibit slower growth due to an increase in primary cilia. However, to our surprise, both KIF24-mutated and KIF24-knockdown (KD) cells grew more vigorously than control cells (Figure 2A, B). These results suggest that KIF24 depletion promotes the growth of Panc1 cells.

**Figure 2.**
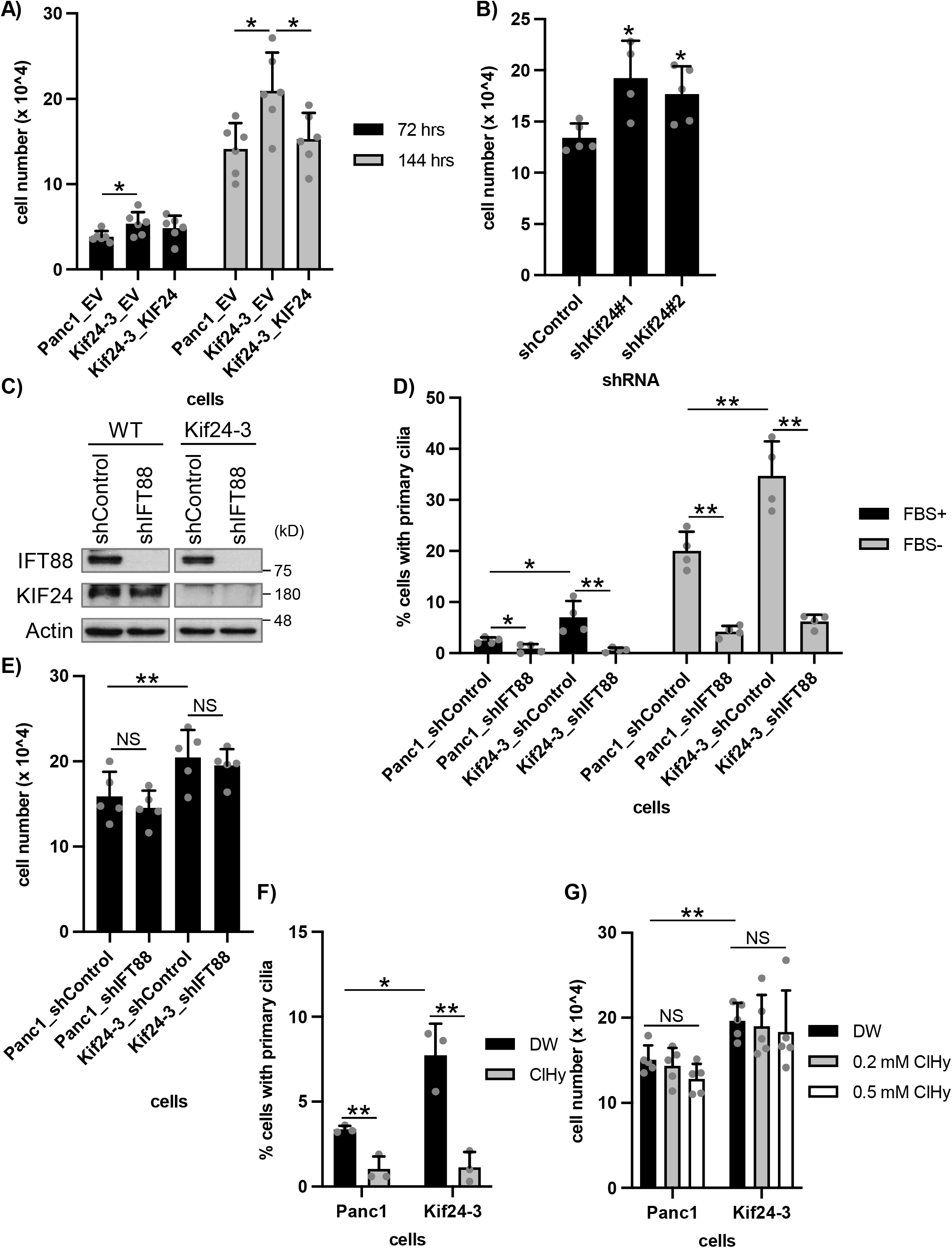
KIF24 depletion enhances proliferation of Panc1 cells *in vitro*. **(A, B)** The indicated Panc1 cells were cultured for 72 or 144 hrs and the number of surviving cells was counted by hemocytometer. The average of six (A) or four to five (B) independent experiments is shown. **(C)** The indicated Panc1 cells were cultured in serum-fed medium for 48 hrs and their extracts were immunoblotted with anti-IFT88 and anti-KIF24-2 antibodies. β-Actin was used as a loading control. **(D)** The indicated Panc1 cells were cultured and immunostained as described in Figure 1B. The percentage of ciliated cells was determined. The average of four independent experiments is shown; >250 cells were scored each time. **(E)** The indicated Panc1 cells were cultured for 144 hrs and the number of surviving cells was counted. The average of five independent experiments is shown. **(F)** The indicated Panc1 cells treated with DW or 0.5 mM ClHy were cultured in serum fed medium for 48 hrs and immunostained as described in Figure 1B. The percentage of ciliated cells was determined. The average of three independent experiments is shown; >250 cells were scored each time. **(G)** The indicated Panc1 cells treated with DW or 0.5 mM ClHy were cultured for 144 hrs and the number of surviving cells was counted. The average of five independent experiments is shown. **(A, B, D-G)** All data are shown as mean ± SD. two-tailed Student’s *t*-test. **, *p* < 0.01; *, *p* < 0.05.

Next, tumor formation in KIF24-mutated Panc1 cells was evaluated in a mouse xenograft model. Kif24-3-tumors were larger than tumors derived from parental cells at early stages (4 and 6 weeks after injection) (Figure S2A). However, the differences in the tumor volume were not significant at later stages (8–14 weeks after injection). Similarly, the excised Kif24-3-tumors weighed moderately heavier than the WT-tumors (14 weeks after injection) (Figure S2B, C). These results suggest that KIF24 depletion tends to accelerate tumor formation of Panc1 cells *in vivo*. Subsequently, the expression of primary cilia was examined in tumor slices. Consistent with previous results of *in vitro* cultivation (Figure 1B, C, 2C, D), more primary cilia were observed in the slices of the Kif24-tumors than in the WT-tumors (Figure S2D, E). Altogether, these results suggest that KIF24 depletion in Panc1 cells induces modest enhancement of tumorigenesis and primary ciliogenesis *in vivo*.

### Primary cilia are irrespective of over-proliferation in KIF24-mutated cells

To test whether primary cilia are related to the hyper-proliferation of KIF24-depleted cells, an Intraflagellar transport (IFT) protein IFT88, which is essential for cilia formation, was stably knocked down (Figure 2C). If forced ciliation by KIF24 loss is linked to the over-proliferation of Panc1 cells, de-ciliation might affect the over-proliferation. In contrast to the substantial decrease in primary ciliation (Figure 2D), cell growth was not affected by silencing of IFT88 in WT and Kif24-3 cells (Figure 2E). To further confirm the primary cilia-independent hyperproliferation of Kif24-3 cells, cells were treated with chloral hydrate (ClHy), which is known to exclude primary cilia from the cell surface (Ho *et al*, 2013; Kobayashi *et al*, 2020). ClHy-treatment significantly reduced ciliation but failed to impact growth in WT and Kif24-3 cells (Figure 2F, G). These results strongly indicate that KIF24 depletion induces over-proliferation of Panc1 cells independent of primary cilium assembly.

### KIF24 depletion induces the centrosome clustering

KIF24 is known to localize to the centrosome not only during interphase but also during mitosis in mammalian cells (Kobayashi *et al*., 2011). We detected centrosomal KIF24 in mitosis of Panc1 cells (Figure S3) and therefore focused on mitotic events in Kif24-3 cells. Immunofluorescence experiments to visualize mitotic morphologies showed that ∼20% of WT (Panc1_EV) cells form multi-polar spindles (Figure 3A, B), which is consistent with a previous study showing that Panc1 cells frequently harbor amplified centrosomes and assemble the multi-polar spindles (Difilippantonio *et al*, 2009). In contrast, the proportion of mitotic cells with multi-polar spindles was significantly decreased in Kif24-3_EV cells compared with Panc1_EV and Kif24-3_KIF24 cells (Figure 3A, B). Next, we stained cells with antibodies recognizing centrin and γ−tubulin to detect centrioles and centrosomes, respectively. Remarkably, this experiment revealed a decrease in multi-polar cells with a concurrent increase in pseudo bi-polar cells in which more than two centrin dots were observed on one side of the spindle (= γ-Tubulin dot(s)) in mitotic Kif24-3 cells (Figure 3C, D), indicating that the supernumerary centrosomes were tightly clustered in KIF24-mutated cells. We also observed more mis-segregated chromosomes in anaphase in Kif24-3 cells (Figure 3E, F), which is typically induced by pseudo bi-polar spindles, supporting that the occurrence of mitotic cells with pseudo bi-polar spindles was increased by KIF24 loss. Moreover, shKif24-Panc1 cells displayed more pseudo bi-polar, rather than multi-polar, spindles (Figure 3G). On the other hand, the percentages of cells with over-amplified centrioles (>4 centrin dots) or fragmented centrosomes (γ-Tubulin foci without centrin) were comparable among analyzed cells (Figure S4A, B), suggesting that KIF24 depletion specifically evokes clustering of centrosomes. CP110 expression was also evaluated because silencing of KIF24 decreases the CP110 protein levels in normal diploid RPE1 cells but not in cancerous U2OS cells (Kobayashi *et al*., 2011). However, CP110 levels were not altered in Kif24-3 cells (Figure S4C), suggesting that, similar to U2OS cells, CP110-dependent centriolar events are unaffected by KIF24 depletion in Panc1 cancer cells. Collectively, these results suggest that a lack of KIF24 induces clustering of centrosomes in Panc1 cells harboring supernumerary centrosomes.

**Figure 3.**
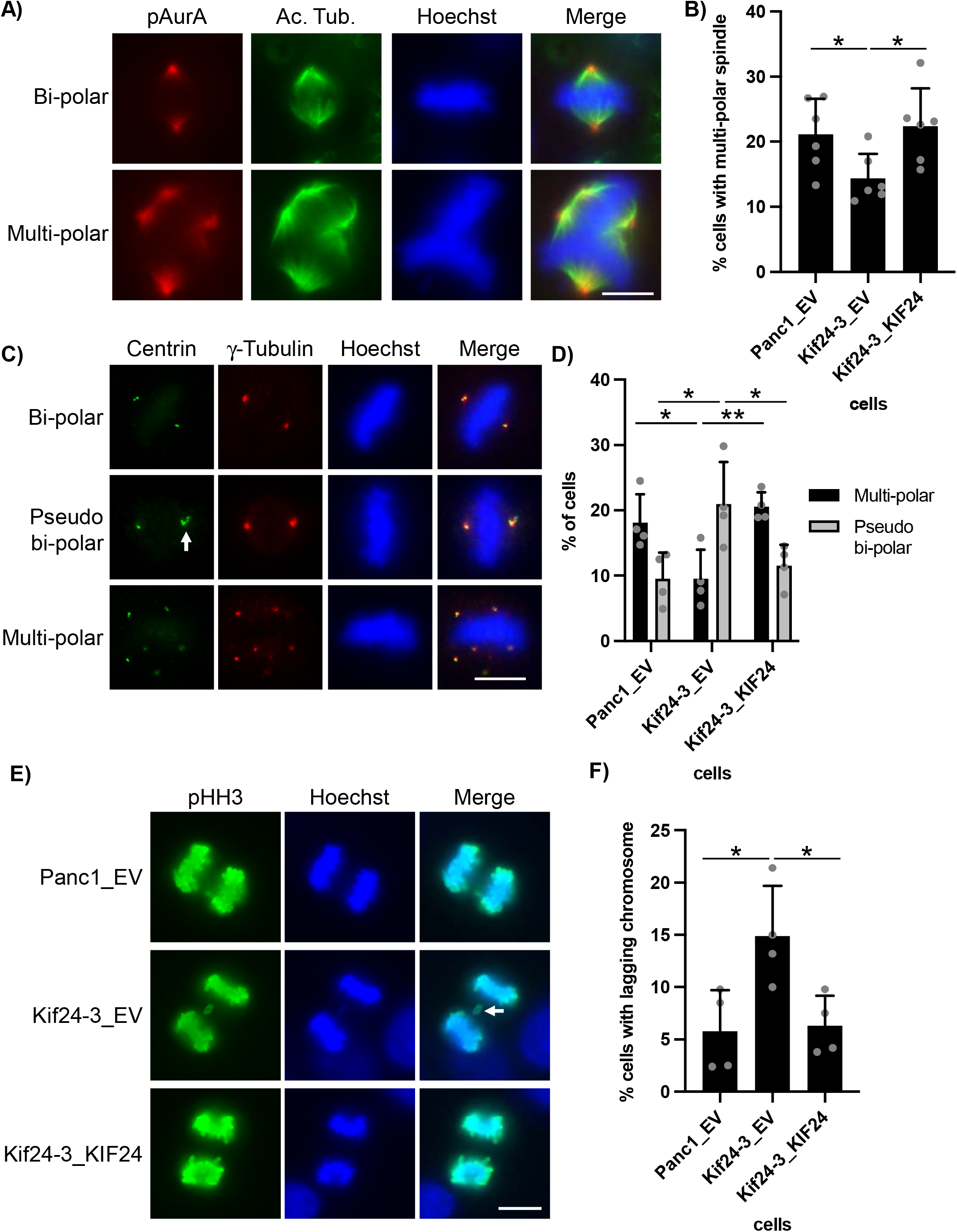
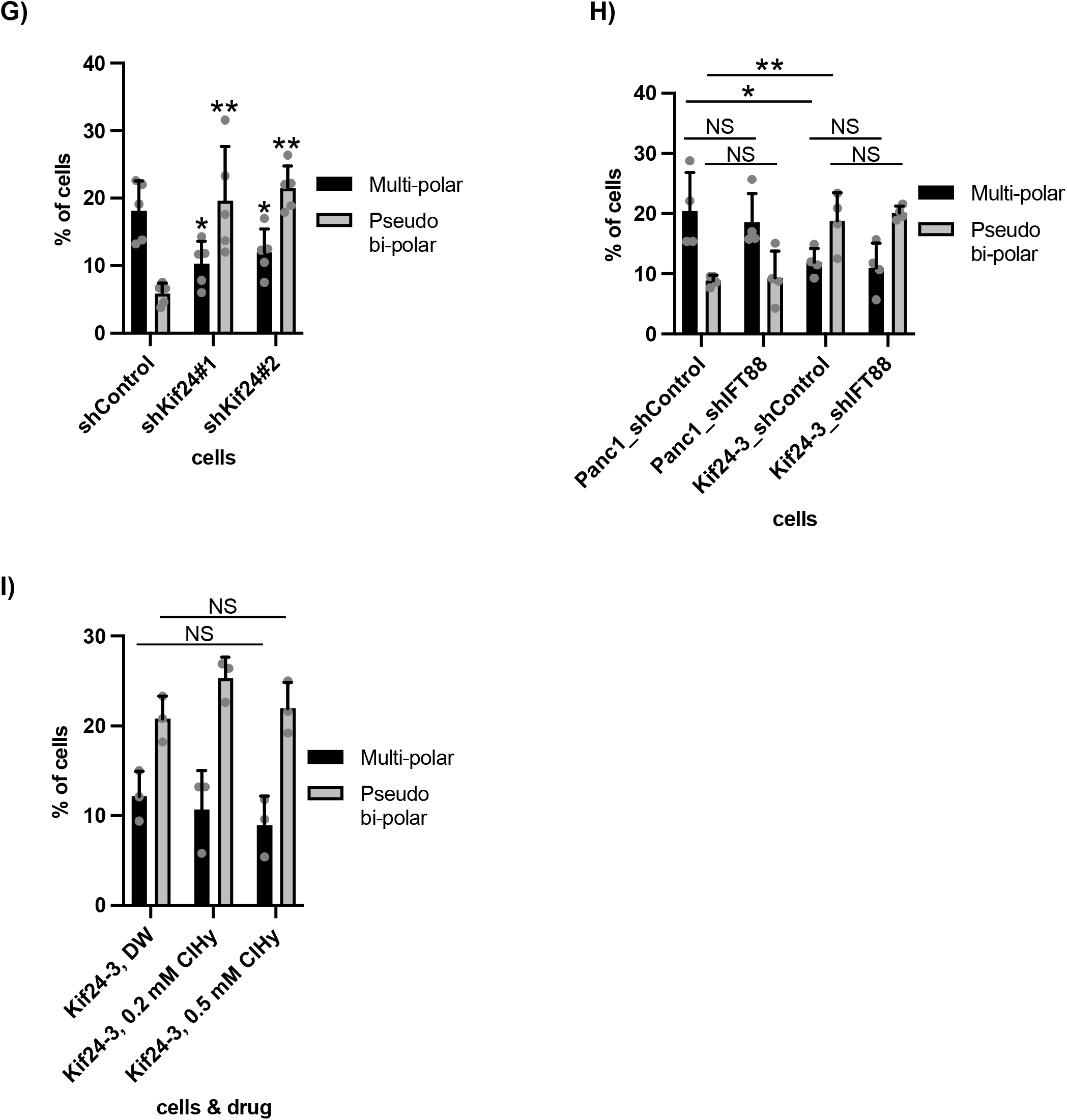
KIF24 depletion induces centrosome clustering in Panc1 cells. **(A, B)** The indicated Panc1 cells were immunostained with anti-Acetylated tubulin (green) and anti-phosphorylated AurA (pAurA) (red) antibodies. (A) DNA was stained with Hoechst (blue). Scale bar, 10 µm. (B) The percentage of cells with multi-polar spindles in metaphase was determined. The average of six independent experiments is shown; >100 cells were scored each time. **(C, D)** The indicated Panc1 cells were immunostained with anti-Centrin (green) and anti-γ-Tubulin (red) antibodies. (C) DNA was stained with Hoechst (blue). Scale bar, 10 µm. (D) The percentage of cells with pseudo bi-polar or multi-polar spindles in metaphase was determined. The average of four independent experiments is shown; >100 cells were scored each time. **(E, F)** The indicated Panc1 cells were immunostained with an anti-phosphorylated Histone H3 (pHH3) (green) antibody. (E) DNA was stained with Hoechst (blue). Scale bar, 10 µm. (F) The percentage of cells with lagging chromosome in anaphase was determined. The average of four independent experiments is shown; >100 cells were scored each time. **(G, H)** The indicated Panc1 cells were immunostained and quantified as described in Figure 3C, D. The average of five (G) or four (H) independent experiments is shown; >100 cells were scored each time. **(I)** Kif24-3 cells treated with DW or ClHy were cultured in serum fed medium for 48 hrs. Cells were immunostained and quantified as described in Figure 3C, D. The average of three independent experiments is shown; >100 cells were scored each time. **(B, D, F-I)** All data are shown as mean ± SD. two-tailed Student’s *t*-test. **, *p* < 0.01; *, *p* < 0.05; NS, no significance.

Subsequently, immunofluorescence studies were conducted in cells treated with shIFT88 or ClHy. Neither IFT88-KD nor ClHy-treatment affected the number of cells with multipolar and pseudo bi-polar spindles in Kif24-3 cells (Figure 3H, I). These data suggest that the centrosome clustering occurs irrespective of primary cilia in KIF24-mutated cells, which is consistent with the over-proliferation of these cells.

### NEK2-mediated phosphorylation and MT-depolymerizing activity are dispensable for the mitotic function of KIF24

The amino acid residues KEC (positions 483–485) are conserved in the Kinesin-13 family of proteins and are important for the MT-depolymerizing activity of KIF24 (Kobayashi *et al*., 2011). In addition, the NEK2-mediated phosphorylation of Thr^622^ and Ser^623^ was found to enhance the MT-depolymerizing activity of KIF24 (Kim *et al*., 2015). KIF24/KEC483-485AAA (KEC)- or KIF24/TS622, 623AA (TS)-expressing Kif24-3 cells were generated to elucidate whether the modification and activation of KIF24 are involved in spindle morphology (Figure 4A). While KIF24/KEC and KIF24/TS failed to suppress the assembly of primary cilia (Figure 4B), these mutants significantly decreased pseudo bi-polar formation and concomitantly increased multi-polar formation in KIF24-depleted cells (Figure 4C). These results suggest that the MT depolymerizing activity and NEK2-mediated phosphorylation of KIF24 are dispensable for its mitotic function, and further support our idea that centrosome clustering occurs independently of primary cilia in KIF24 depleted cells.

**Figure 4.**
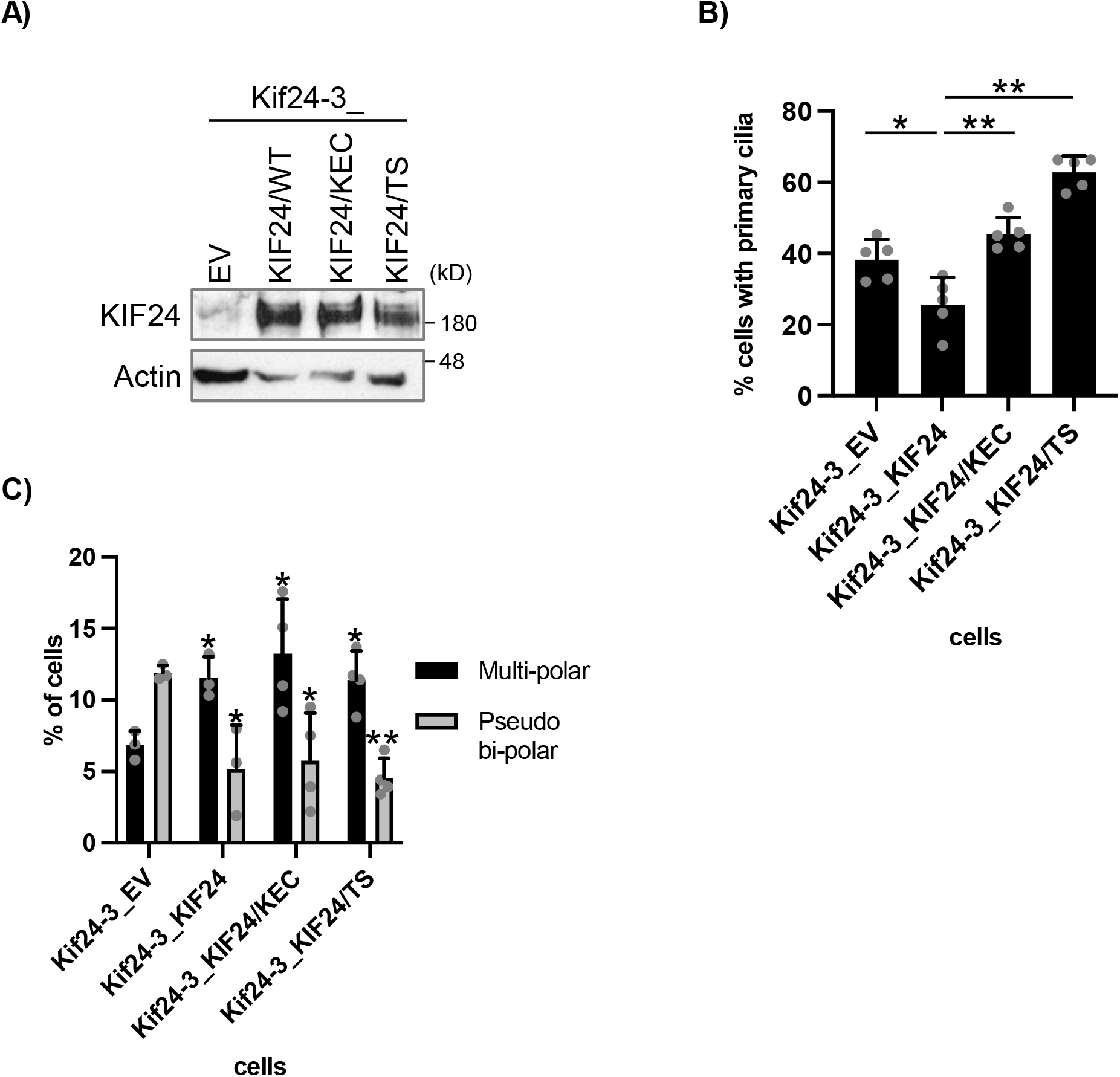
NEK2-phosphorylation or MT-depolymerizing activity dead mutants are sufficient for mitotic function of KIF24. **(A)** The indicated Panc1 cells were cultured and immunoblotted as described in Figure 1A. **(B)** The indicated Panc1 cells were cultured and immunostained as described in Figure 1B. The percentage of ciliated cells was determined. The average of five independent experiments is shown; >250 cells were scored each time. **(C)** The indicated Panc1 cells were cultured, immunostained, and quantified as described in Figure 3C, D. The average of three to four independent experiments is shown; >100 cells were scored each time. **(B, C)** All data are shown as mean ± SD. two-tailed Student’s *t*-test. **, *p* < 0.01; *, *p* < 0.05

### HSET/KIFC1 inhibition suppresses centrosome clustering caused by KIF24 depletion

The mitotic kinesin, HSET/KIFC1, promotes centrosome clustering (Kleylein-Sohn *et al*, 2012), and treatment with the allosteric HSET inhibitor, CW069, provokes multi-polar spindles in centrosome-amplified cells (Watts *et al*, 2013). These findings prompted us to examine whether HSET inhibition suppresses pseudo bi-polar formation in KIF24-depleted cells. CW069 treatment clearly induced formation of multi-polar spindles instead of pseudo bi-polar assemblies in Kif24-3 cells, and the ratios were comparable in Panc1_EV and Kif24-3_KIF24 cells (Figure 5A). These results suggest that KIF24 functions upstream of HSET in the centrosome clustering cascade. On the other hand, HSET on the mitotic spindle microtubules was unaffected in KIF24-depleted cells (Figure 5B). KIF24 at spindles also remained unchanged by CW069 treatment (Figure 5C). These results suggest that KIF24 is unrelated to HSET localization and *vice versa*, in mitosis.

**Figure 5.**
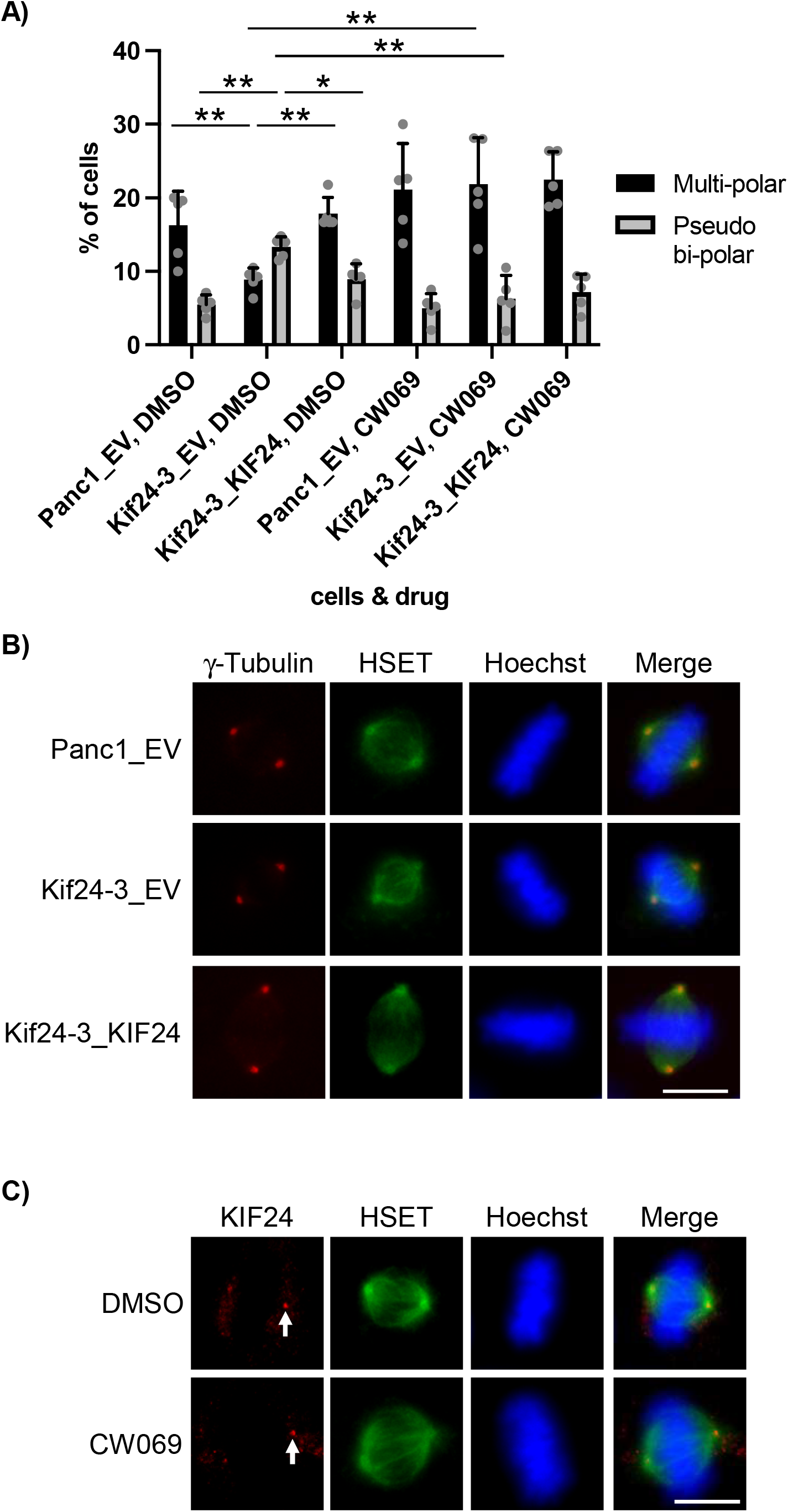
HSET/KIFC1 inhibition suppresses centrosome clustering induced by KIF24 depletion. **(A)** The indicated Panc1 cells treated with DMSO or 50 µM CW069 for 4 hrs were cultured, immunostained, and quantified as described in Figure 3C, D. The average of five independent experiments is shown; >100 cells were scored each time. **(B)** The indicated Panc1 cells were immunostained with anti-HSET/KIFC1 (green) and anti-γ-Tubulin (red) antibodies. DNA was stained with Hoechst (blue). Scale bar, 10 µm. **(C)** Panc1 cells treated with DMSO or 50 µM CW069 for 4 hrs were immunostained with anti-HSET/KIFC1 (green) and anti-KIF24 (red) antibodies. Arrows indicate KIF24 at spindles. DNA was stained with Hoechst (blue). Scale bar, 10 µm. **(A)** All data are shown as mean ± SD. two-tailed Student’s *t*-test. **, *p* < 0.01; *, *p* < 0.05

### KIF24 depletion ameliorates mitotic incapability in Panc1 cells

Live-cell imaging of mitotic progression was subsequently performed in KIF24-mutated cells. To visualize the chromosome dynamics during mitosis, histone H2B-mCherry stably expressing Panc1 or Kif24-3 cells were generated. Time-lapse imaging revealed that the mitotic duration from nuclear envelope breakdown (NEB) to anaphase onset was slightly but significantly shortened in Kif24-3 cells (Figure 6A, B). Moreover, the ratio of cells that entered mitosis but failed to divide was largely decreased in Kif24-3 cells (Figure 6A, C). These results suggest that KIF24 depletion improves the mitotic progression in centrosome-amplified Panc1 cells, probably leading to their accelerated proliferation.

**Figure 6.**
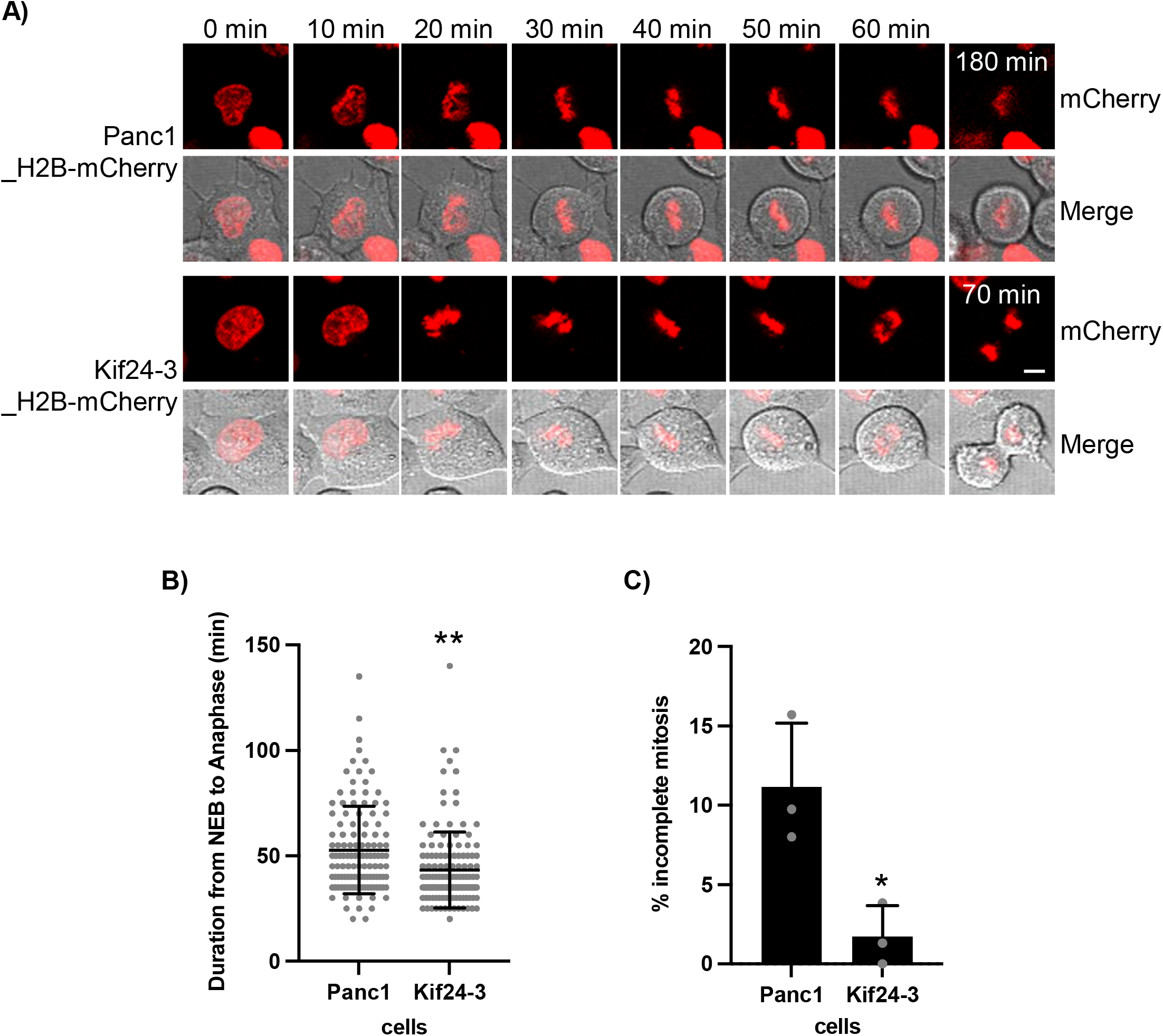
KIF24 depletion restores impaired mitotic events in Panc1 cells. **(A)** Frames from live cell imaging of Panc1 or Kif24-3 cells stably expressing H2B-mCherry. Scale bar, 10 µm. **(B)** Time from nuclear envelope breakdown (NEB) to anaphase onset in indicated cells was measured. n = 119 (Panc1_H2B-mCherry), 135 (Kif24-3_H2B-mCherry). **(C)** The percentage of cells that entered and stayed >180 min in mitosis was determined. The average of three independent experiments is shown; >40 cells were scored each time. **(B, C)** All data are shown as mean ± SD. two-tailed Student’s *t*-test. **, *p* < 0.01; *, *p* < 0.05

### KIF24 depletion suppresses multipolar spindle formation and enhances cell growth specifically in centrosome-amplified PDAC cells

To test whether KIF24 controls centrosome clustering in other PDAC cells, shKIF24-expressed MiaPaCa2 or Hs766t cells were generated (Figure S1C). A previous report indicated that centrosomes are amplified in Hs766t cells but not in MiaPaCa2 cells (Difilippantonio *et al*., 2009). These PDAC cells rarely assembled primary cilia even after KIF24 silencing, suggesting that these cells near-completely lost the ability to ciliate (Figure 7A). In MiaPaCa2 cells, both control and KIF24-depleted cells assembled multipolar spindles with only ∼5% frequency (Figure 7B, C). In contrast, multipolar spindles were detected in ∼27% of control Hs766t cells, and this frequency was considerably reduced by KIF24 depletion (Figure 7B, C). Concomitantly, KIF24-depleted Hs766t cells grew more vigorously than control Hs766t cells, in sharp contrast with MiaPaCa2 cells (Figure 7D). These results suggest that KIF24 induces multipolar spindle formation and thereby slow growth, specifically in centrosome-amplified PDAC cells.

**Figure 7.**
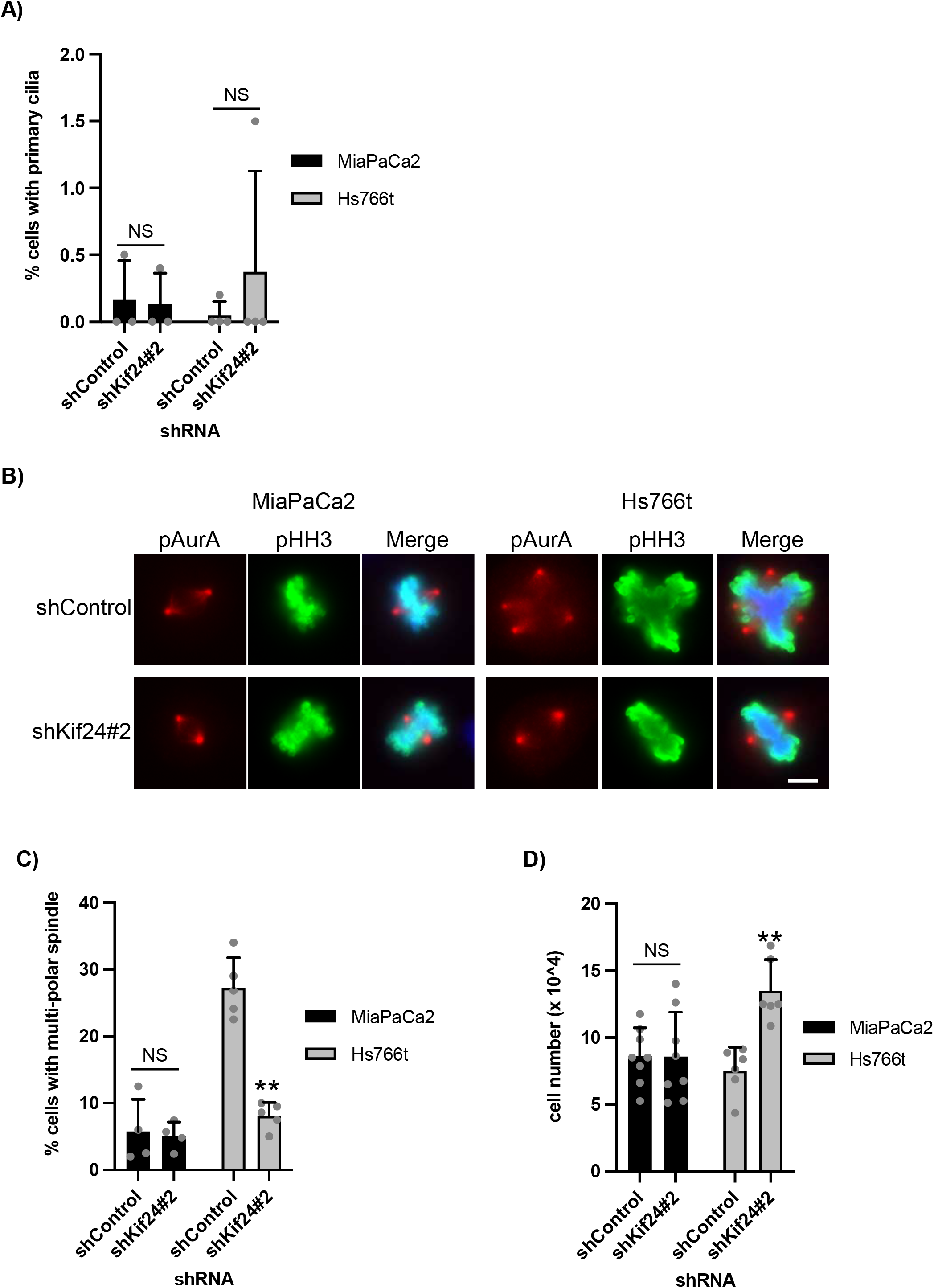
KIF24 depletion suppresses multi-polar spindle formation and promotes cell growth in centrosome-amplified PDAC cells. **(A)** The indicated PDAC cells were cultured and immunostained as described in Figure 1B. The percentage of ciliated cells was determined. The average of three (MiaPaCa2) or four (Hs766t) independent experiments is shown; >250 cells were scored each time. **(B, C)** The indicated PDAC cells were immunostained with anti-pHH3 (green) and anti-pAurA (red) antibodies. (B) DNA was stained with Hoechst (blue). Scale bar, 10 µm. (C) The percentage of cells with multi-polar spindles in metaphase was determined. The average of four (MiaPaCa2) or five (Hs766t) independent experiments is shown; >100 cells were scored each time. **(D)** The indicated cells were cultured for 96 hrs (MiaPaCa2) or 192 hrs (Hs766t) and the number of surviving cells was counted by hemocytometer. The average of eight (MiaPaCa2) or six (Hs766t) independent experiments is shown. **(A, C, D)** All data are shown as mean ± SD. two-tailed Student’s *t*-test. **, *p* < 0.01; *, *p* < 0.05; NS, no significance.

## Discussion

This study analyzed PDAC cells with supernumerary centrosomes revealing that KIF24 suppresses the clustering of excess centrosomes during mitosis as well as the assembly of primary cilia. NEK2-mediated phosphorylation and MT-depolymerizing activity were dispensable for the mitotic role of KIF24, unlike the ciliary role, indicating that distinct regulatory systems govern the dual functions of KIF24 (Figure S5). These mitotic and ciliary phenotypes in cells depleted of KIF24 are thought to exert reciprocal effects on cell proliferation. KIF24-depleted PDAC cells overgrew *in vitro*, in which primary cilia are rarely assembled (∼10% in Kif24-3 cells, Figure 1C), probably due to improved mitosis. On the other hand, the tumors derived from KIF24-depleted cells were not significantly larger than those derived from parental cells in the mouse xenograft model. As more primary cilia were assembled in the tumors than in the cultured cells (∼25% in Kif24-3 tumor slices, Figure S2E), it is plausible that these organelles opposed the *in vivo* growth of Kif24-3 cells. Alternatively, aneuploidy provoked by chromosome mis-segregation in Kif24-3 cells may result in heterogeneous populations of cells with diverse proliferative profiles during long-term tumor growth. This notion is supported by data showing that Kif24-3 tumors exhibited larger variation than wild-type tumors. On the other hand, as the transient assembly of primary cilia can promote cell proliferation by activating the hedgehog (Hh) pathway in medulloblastoma cells (Ho *et al*, 2020), we cannot exclude the possibility that primary cilia induced by KIF24 loss positively affects tumor growth *in vivo*, where implanted tissue is presumably exposed to Hh ligands. A clinical report showing that expression of primary cilia is correlated with poor prognosis of PDAC patients also suggests a growth-promoting of primary cilia in PDAC *in vivo* (Emoto *et al*, 2014). Several MT-binding proteins are known to be involved in the regulation of multipolar spindle formation in cancer cells but not in PDAC cells. HSET promotes the clustering of amplified centrosomes in breast cancer and melanoma cells (Kleylein-Sohn *et al*., 2012). HSET interacts with a centrosomal protein CEP215 during its operation (Chavali *et al*, 2016). IFT-B proteins, which are essential for cilia assembly, are also associated with HSET to promote centrosome clustering in several centrosome-amplified cells (Vitre *et al*, 2020), whereas we did not detect any alterations of centrosome clustering in IFT88-depleted Panc1 cells. Contrary to CEP215 and IFT-B proteins, KIF24 is unlikely to directly associate with HSET but probably acts upstream of HSET in PDAC cells based on our data. Another kinesin, KIF18A, facilitates the proliferation of colorectal and breast cancer cells through the regulation of centrosome fragment formation (Marquis *et al*, 2021). The association of a centrosomal protein, CPAP, with MT is required for centrosome clustering in breast and lung cancer cells (Mariappan *et al*, 2019). CPAP antagonizes CP110 in centriole elongation and is required for ciliogenesis (Schmidt *et al*, 2009; Wu & Tang, 2012). Given that KIF24 interacts and cooperates with CP110 in the suppression of primary cilia formation (Kobayashi *et al*., 2011), it is conceivable that KIF24 prevents the CPAP-MT association and thereby induces nucleation of multipolar spindles.

Although we showed that primary cilium assembly has no bearing on centrosome clustering in PDAC cells, the possibility that primary cilia or their remnants after resorption influence mitosis cannot be excluded. Pharmacological inhibition of mitotic kinases, Aurora A or PLK1, induces mitotic primary cilia in normal mouse IMCD3 cells (Bowler *et al*, 2019). It was recently reported that NEK2 knockout causes the expression of primary cilia or their remnants in mitotic RPE1 cells (Viol *et al*, 2020). As described above, NEK2 phosphorylates and activates KIF24 to disassemble primary cilia in G2–M phase (Kim *et al*., 2015). Indeed, mitotic primary cilia in KIF24-mutated Panc1 cells were found; however, we did not observe detectable alterations in centrosome clustering (data not shown). It will be interesting to determine whether and how primary cilia influence various mitotic events in cancer cells.

Centrosome clustering is an attractive target for cancer therapy because it is frequently observed in cancer cells but not in normal cells. Based on our study, potentiation of the mitotic activity of KIF24 could be a valid intervention strategy for PDAC, although we need to consider that loss of primary cilia by KIF24 might aggravate this cancer at present. Future studies clarifying the mechanistic details of KIF24 in spindle formation will represent a promising therapeutic target to develop novel centrosome de-clustering drugs for PDAC.

## Materials & Methods

### Cell culture

Panc1 (American Type Culture Collection), MiaPaCa2, Hs766t (Li *et al*, 2013), Kif24-3 Panc1 (this study), and Lenti-X 293T (gift from M. Hagiwara) cells were grown in Dulbecco’s Modified Eagle Medium (DMEM) (Nacalai tesque) supplemented with 10% Fetal Bovine Serum (FBS) (Biosera) and 100 units/ml penicillin and 100 μg/ml streptomycin (P/S) (Nacalai tesque).

### Antibodies and Reagents

Antibodies used in this study include mouse anti-glutamylated tubulin (GT335) (1:1000 (IF), Adipogen, AG-20B-0020), rabbit anti-ARL13B (1:1000 (IF), Proteintech, 17711-1-AP), mouse anti-ARL13B (1:1000 (IF), NeuroMab, 75-287), rabbit anti-KIF24 (1:200 (IF) (Kobayashi *et al*., 2011)), rabbit anti-KIF24-2 (1:500 (WB), this work), rabbit anti-CP110 (1:1000 (WB) (Chen *et al*, 2002)), mouse anti-phosphorylated AurA (1:100 (IF), Cell signaling, #3079), mouse anti-acetylated tubulin (1:1000 (IF), SIGMA, T7451), goat anti-γ-Tubulin (1:400 (IF), Santa Cruz, sc-7396), mouse anti-centrin (1:1000 (IF), Millipore, 04-1624), mouse anti-phosphorylated Histone H3 (1:1000 (IF), MAB institute, MABI0012), rabbit anti-IFT88 (1:1000 (WB), Proteintech, 13967-1-AP), mouse anti-KIFC1/HSET (1:500 (IF), Santa Cruz, sc-100947), and mouse anti-β-Actin (1:1000, Santa Cruz, sc-47778). A rabbit anti-KIF24-2 antibody was produced by immunizing a glutathione-S-transferase (GST) fusion protein containing residues 1201-1368 of KIF24 into rabbits and purified as described previously (Kobayashi *et al*., 2011). Reagents used in this study include Chloral Hydrate (ClHy) (Nacalai tesque, 07922-62), Thymidine (Sigma Aldrich, 89270), CW069 (Selleckchem, S7336), and Hoechst 33342 (Nacalai tesque, 04915-82).

### Plasmids

To generate guide RNA (gRNA) targeting KIF24, annealed oligo was inserted into pSpCas9(BB)-2A-Puro (PX459) V2.0 (Addgene) (Ran *et al*, 2013). To generate short hairpin RNA (shRNA) targeting IFT88, KIF24, or negative control annealed oligo was inserted into pLKO.1 (Addgene) (Stewart *et al*, 2003). Oligo DNAs are listed in Table S1. To generate KIF24 wild-type or KEC483-485AAA (KEC), human KIF24 fragments encoding residue 1-4383 was excised from pEGFP-C1-KIF24 (Kobayashi *et al*., 2011) and sub-cloned into pLVX-IRES-Puro (Clontech). KIF24/TS621-622AA (TS) construct was made by PCR-based mutagenesis using primers listed in Table S1. To generate H2B-mCherry, human H2BC11 fragment encoding residue 1-381 was amplified by PCR using primers listed in Table S1, and sub-cloned into pLVX-mCherry-IRES-Puro. pLVX-mCherry-IRES-Puro was constructed by replacing GFP of pLVX-GFP-IRES-Puro with mCherry (Kim *et al*., 2015). Plasmid transfection into Panc1 and Lenti-X 293T cells was performed using Lipofectamine 2000 (Invitrogen) and PEI Max (Polysciences) according to the manufacturer’s instruction, respectively.

### Generation of Kif24-3 cells

PX459-KIF24 plasmid was transfected into Panc1 cells using Lipofectamine 2000. Transfected cells were cultured in medium with 5 µg/ml puromycin (Nacalai tesque) for 72 hrs and singly plated into 96-well plates. Genome DNA was extracted from survival cells using QuickExtract DNA Solution 1.0 (Epicentre), and amplified PCR products using primers listed in Table S1 were sub-cloned into pGEM-T Easy (Promega). Purified plasmid DNAs were sequenced using M13 primers.

### Generation of stable cells

Lentivirus supernatant was produced by co-transfection of pLVX-IRES-Puro (Empty Vector (EV)), pLVX-KIF24-IRES-Puro (KIF24), pLVX-KIF24/TS-IRES-Puro (KIF24/TS), pLVX-KIF24/KEC-IRES-Puro (KIF24/KEC), pLVX-H2B-mCherry-IRES-Puro (H2B-mCherry), pLKO.1-shControl, pLKO.1-shKif24#1, pLKO.1-shKif24#2, or pLKO.1-shIFT88 with Δ8.9, pcRev, and VSVG plasmids (gift from M. Hagiwara) into Lenti-X 293T cells using PEI Max. The virus supernatant was harvested 72 hrs post-transfection and concentrated using Lenti-X Concentrator (Clontech). Panc1, MiaPaCa2, and Hs766t cells were incubated with virus in the presence of 5 µg/ml polybrene (Nacalai tesque) for 72 hrs. The infected cells were subsequently cultured in medium with 3 µg/ml (Panc1), 0.5 µg/ml (MiaPaCa2), or 1.5 µg/ml (Hs766t) puromycin for 8 to 20 days. Established cells were cultured in medium with puromycin. Panc1/WT_EV cells were generated previously (Kobayashi *et al*., 2020).

### Cell growth assay

1 × 10^4^ cells were seeded in 24-well plate and cultured for 72 or 144 hrs (Panc1), 96 hrs (MiaPaCa2), or 192 hrs (Hs766t). Number of trypsinized cells was counted using hemocytometer.

### Western blotting

Cells were lysed with lysis buffer (50 mM Hepes-NaOH pH 7.5, 150 mM NaCl, 5 mM EDTA, 0.5% NP-40, 10% Glycerol, 1 mM DTT, 0.5 mM PMSF, 2 µg/ml leupeptin, 5 mM NaF, 10 mM β-Glycerophosphate, and 1 mM Na_3_VO_4_) at 4 °C for 30 min. A 20 µg lysate was loaded and analyzed using SDS-PAGE and immunoblotting.

### Immunofluorescence microscopy

Cultured cells were fixed with cold Methanol for 5 min, 4% Paraformaldehyde (Nacalai tesque)/PBS for 10 min, or 3.7% Formalin (Nacalai tesque)/PBS for 10 min. After permeabilization with 0.2% Triton X-100/PBS for 10 min, slides were blocked with 5% BSA/PBS prior to incubation with primary antibodies. Primary and secondary antibodies were diluted to the desire concentrations using 5% BSA/PBS. Secondary antibodies used were AlexaFluor488- or AlexaFluor594-conjugated donkey anti-mouse, anti-rabbit, or anti-goat IgG (Invitrogen). Cells were stained with Hoechst33342 to visualize DNA. Mounted slides with PermaFluor Mounting Medium (Thermo Scientific) were observed and imaged using AxioObserver with a 63× lens. Tumors were fixed with 3.7% Formalin/PBS at 4 °C for 12 hrs, sequentially equilibrated with 10%, 20%, and 30% Sucrose/PBS, and embedded with OCT compound (Sakura Finetek) at -80 °C. The frozen tumors were sliced into 10 µm-thick sections using Cryostat NX70 (Thermo) and mounted on MAS-coated slide (Matsunami). The mounted sections were fixed with Acetone for 15 min, soaked into boiled water for 15 min to retrieve antigens, and permeabilized with 0.2% Triton X-100/PBS for 10 min. After the permeabilization, procedures were same as cultured cells.

### Quantitative PCR (qPCR)

Total RNA was isolated from cultured cells using Sepasol (Nacalai tesque), and following reverse transcription reaction was performed using ReverTra Ace qPCR RT kit (TOYOBO). qPCR was performed using THUNDERBIRD SYBR qPCR mix (TOYOBO) and LightCycler96 (Roche). All reaction was conducted according to the manufacturer’s instruction. Primers are listed in Table S1.

### Live Imaging

H2B-mCherry-expressed Panc1 or Kif24-3 cells were seeded onto a Cellview cell culture dish (Greiner) and treated with 2 mM Thymidine for 40 hrs. 9 hrs after Thymidine wash-out, cells were imaged using LSM710 (Zeiss) operated by Zen software (Zeiss). 6 Z-stack images were acquired every 5 min for 8 hrs using 20 x objective lens (Zeiss).

### Xenograft

2 × 10^6^ Panc1 cells in PBS were subcutaneously injected into 6-weeks-old female nude mouse (Nihon SLC). Tumors were measured using a caliper weekly, and their volumes were calculated using the formula: length x width x height x 0.5. After 14-weeks, tumors were excised and weighed.

### Statistical Analysis

The statistical significance of the difference was determined using two-tailed Student’s *t*-test. Figure legends indicate the number of independent replicates conducted and the number of cells analyzed for each replicate. Differences were considered significant when *p* < 0.05.

## Acknowledgements

We thank M. Hagiwara (Kyoto University) for providing Lenti-X 293T cells, and Δ8.9, pcRev, and VSVG plasmids.

T.K. was supported by grants from JSPS KAKENHI (15H01215, 15K07931, 18K06627, 21K06528), The Kurata Memorial Hitachi Science and Technology Foundation, Takeda Science Foundation, Daiichi Sankyo Foundation of Life Science, Sagawa Foundation for Promotion of Cancer Research, Mochida Memorial Foundation for Medical and Pharmaceutical Research, and Foundation for Nara Institute of Science and Technology.

## Competing interests

The authors declare no competing financial interests.

## Notes

### Competing Interest Statement

The authors have declared no competing interest.

