## Supplemental Figures for "KIF24 controls the clustering of supernumerary centrosomes in pancreatic ductal adenocarcinoma cells"

A)

**Kif24-3 (exon 1)**

```
WT        TCAGTATCCCAGA GCGTCATCTTCAGACAAGCAGCCTGCGCATC
Allele A  TCAGTATCCCAGAAGCGTCATCTTCAGACAAGCAGCCTGCGCATC
Allele B  TC----------- --------------------------------
Allele C  TCAGT-------- GCGTCATCTTCAGACAAGCAGCCTGCGCATC
Allele D  TCAGTATCCCAGA ---TCATCTTCAGACAAGCAGCCTGCGCATC

WT        AAATCTCAGGAATTAAGATCTGGCCCTCGCAGACA
Allele A  AAATCTCAGGAATTAAGATCTGGCCCTCGCAGACA  1 bp insertion
Allele B  -----------------------------G----CA  74 bp deletion
Allele C  AAATCTCAGGAATTAAGATCTGGCCCTCGCAGACA  8 bp deletion
Allele D  AAATCTCAGGAATTAAGATCTGGCCCTCGCAGACA  3 bp deletion
```

WT 1368 a.a.
Allele A 100 (75 + 25) a.a.
Allele B 75 (71 + 4) a.a.
Allele C 97 (72 + 25) a.a.
Allele D 1367 (1366 + 1) a.a. (75$^{th}$ ER → D)

B)

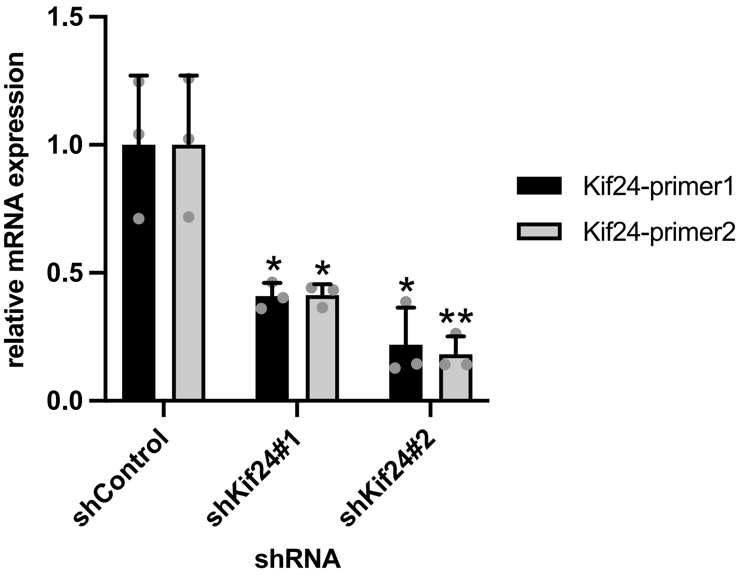


C)

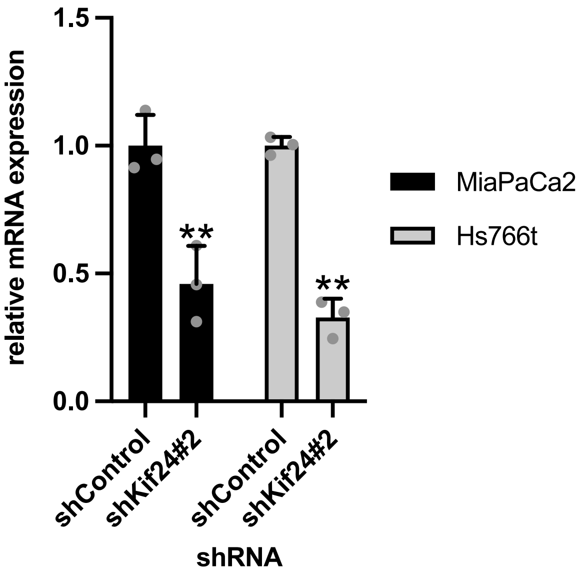


**Figure S1**

**Mutations in Kif24-3 Panc1 cells and knock-down efficiency of shKIF24-expressed PDAC cells**

**(A)** (upper) Four different mutations (alleles A, B, C, and D) of the KIF24 gene in Kif24-3 Panc1 cells (note that Panc1 cells are frequently multiploid). Red sequences represent the target of guide RNAs with underlined protospacer adjacent motif (PAM). (lower) Proteins translated from each clone. Green shows incorrect residues. **(B, C)** shKif24-expressed Panc1 cells (B) or MiaPaCa2 and Hs766t cells (C) were cultured in serum-fed medium for 48 hrs. Relative amount of KIF24 mRNA was determined using quantitative PCR. GAPDH was used as a control. Average of three independent experiments is shown. (C) Kif24-primer2 pairs were used.

**(B, C)** All data are shown as mean ± SD. two-tailed Student's *t*-test. **, $p < 0.01$; *, $p < 0.05$.

**Figure S1**

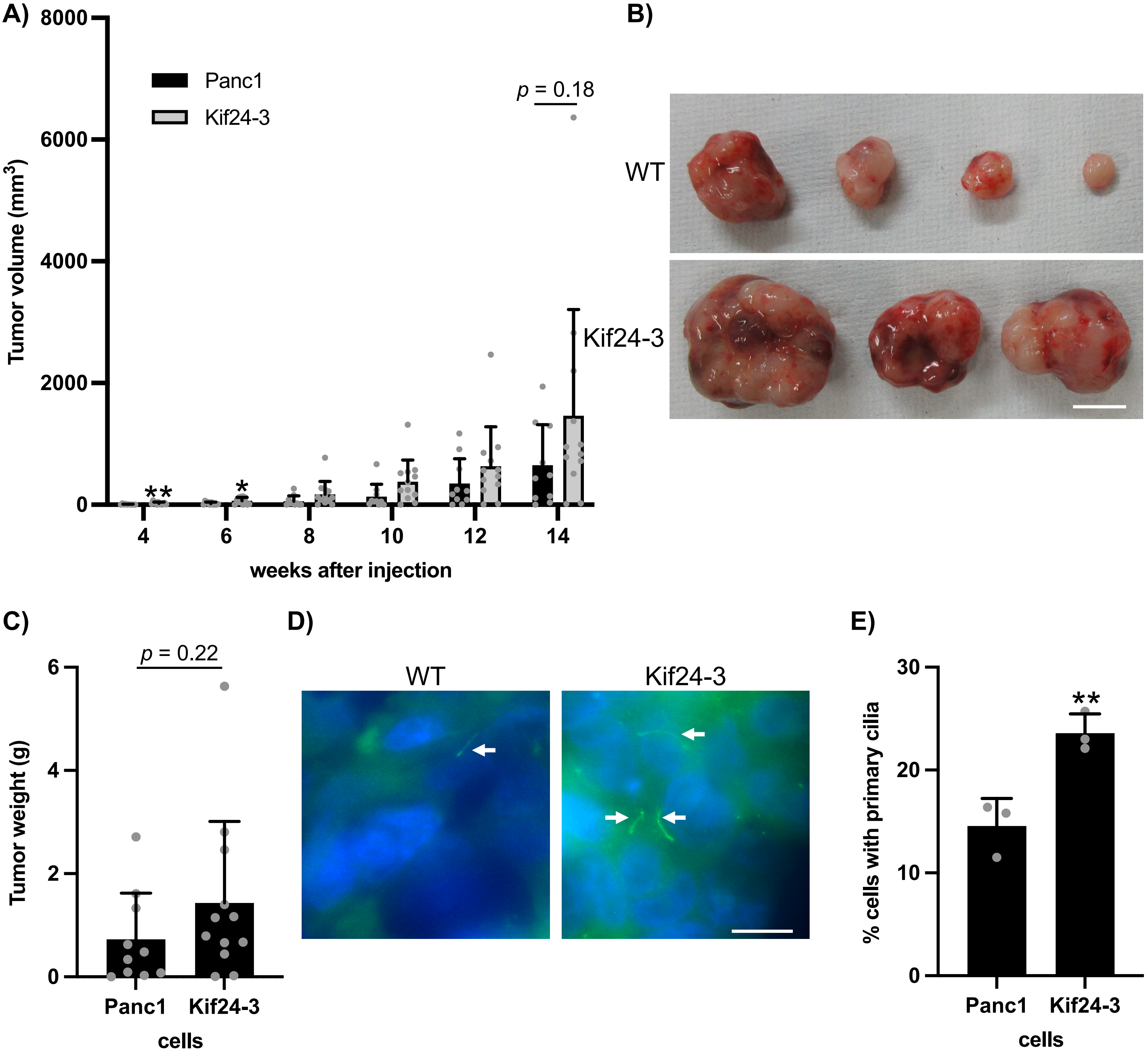


**Figure S2**

**KIF24 mutation moderately promotes tumor formation *in vivo***

**(A-C)** The indicated Panc1 cells were injected to nude mouse. n = 10 (WT), 12 (Kif24-3). (A) The volume of tumor was measured every 2 weeks. (B) Excised tumors were imaged. Scale bar, 10 mm. (C) The weight of excised tumors was measured. **(D, E)** Tumor slices were immunostained as described in Figure 1B. (D) Arrows indicate primary cilia. DNA was stained with Hoechst (blue). Scale bar, 10 µm. (E) The percentage of ciliated cells was determined. The average of three independent experiments is shown; >250 cells were scored each time.

**(A, C, E)** All data are shown as mean ± SD. two-tailed Student's *t*-test. **, $p < 0.01$; *, $p < 0.05$.

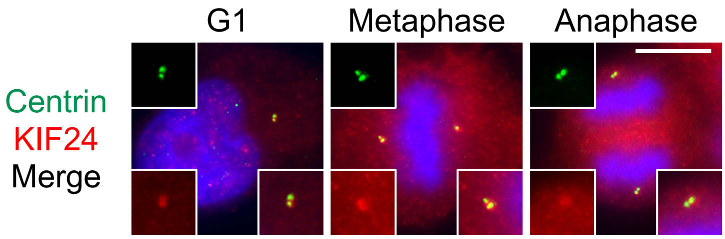


**Figure S3**
**KIF24 localizes to centrosomes in Panc1 cells**
Panc1 cells were immunostained with anti-Centrin (green) and anti-KIF24 (red) antibodies. DNA was stained with Hoechst (blue). Scale bar, 10 µm.

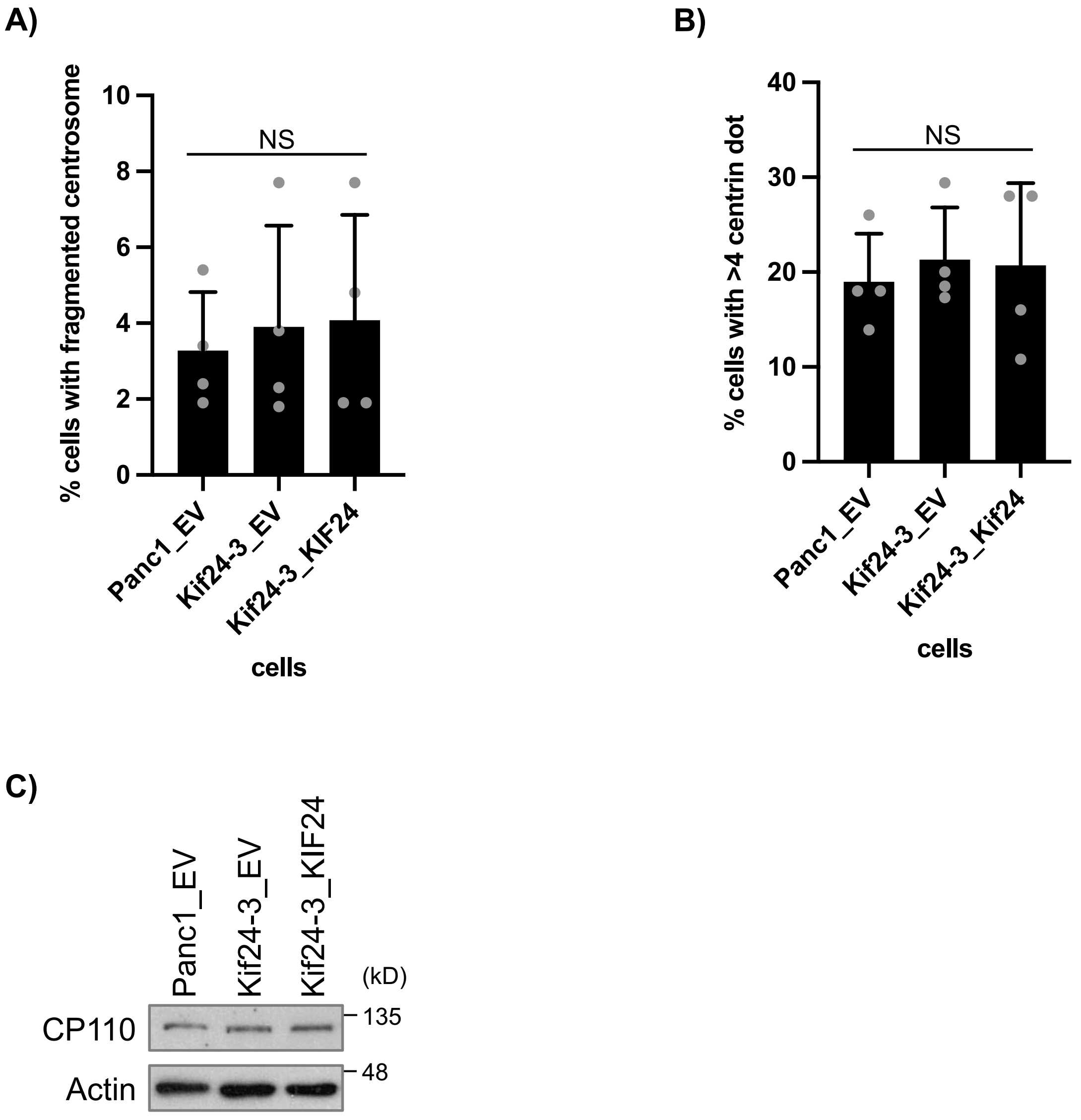


**Figure S4**

**KIF24 depletion does not impinge on centriole duplication and centrosome fragmentation in Panc1 cells**

**(A, B)** The indicated Panc1 cells were immunostained as described in Figure 5C. (A) The percentage of cells with γ-Tubulin-positive and Centrin-negative dot in metaphase was determined. The average of four independent experiments is shown; >100 cells were scored each time. (B) The percentage of cells with >4 centrioles in metaphase was determined. The average of four independent experiments is shown; >100 cells were scored each time. **(C)** The indicated Panc1 cells were cultured in serum-fed medium for 48 hrs. Cell extracts were immunoblotted with an anti-CP110 antibody. b-Actin was used as a loading control. **(A, B)** All data are shown as mean ± SD. two-tailed Student's *t*-test. NS, no significance.

**Figure S4**

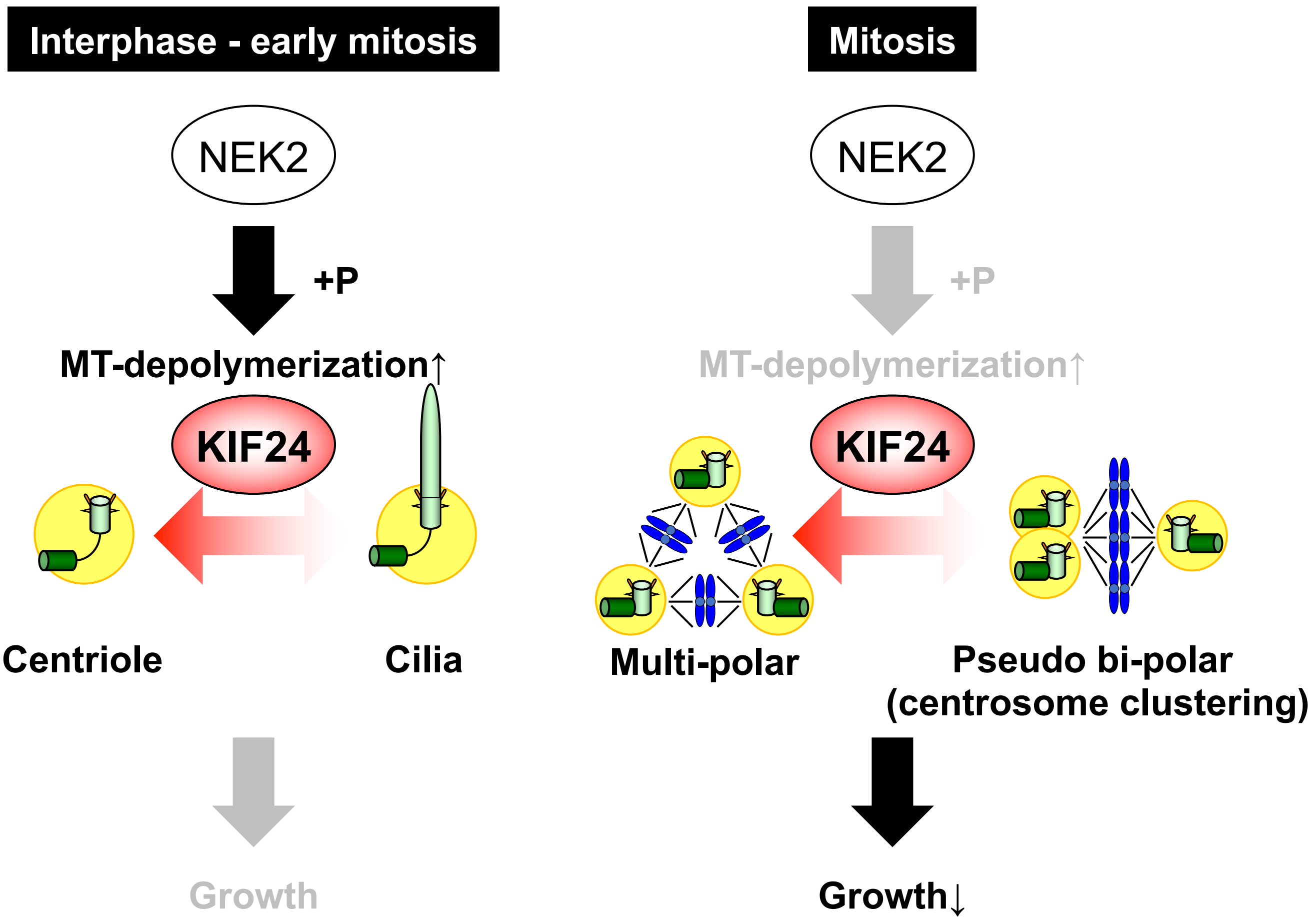


**Figure S5**
**Roles of KIF24 in centrosome-amplified PDAC cells**

| Name | Sequence (5' to 3') | use |
|---|---|---|
| hKif24KO_1_F | CACCGTTGTCTGAAGATGACGCTCT | Guide RNA |
| hKif24KO_1_R | AAACAGAGCGTCATCTTCAGACAAC | Guide RNA |
| hKif24_1_check_F | ATGGCATCCTGGTTATATGAATG | Genome PCR |
| hKif24_1_check_R | AATCCCCCAGTATTGCAGAAAGG | Genome PCR |
| shControl_F | CCGG GCAATCGAAGCTCGGCTACAT CTCGAG ATGTAGCCGAGCTTCGATTGC TTTTTG | shRNA |
| shControl_R | AATTCAAAAA GCAATCGAAGCTCGGCTACAT CTCGAG ATGTAGCCGAGCTTCGATTGC | shRNA |
| shKif24#1_F | CCGG AGGAACACCCTGGAGAATAGC CTCGAG GCTATTCTCCAGGGTGTTCCT TTTTTG | shRNA |
| shKif24#1_R | AATTCAAAAA AGGAACACCCTGGAGAATAGC CTCGAG GCTATTCTCCAGGGTGTTCCT | shRNA |
| shKif24#2_F | CCGG GGGAAGAAAGCTCCGAAATAG CTCGAG CTATTTCGGAGCTTTCTTCCC TTTTTG | shRNA |
| shKif24#2_R | AATTCAAAAA GGGAAGAAAGCTCCGAAATAG CTCGAG CTATTTCGGAGCTTTCTTCCC | shRNA |
| shIFT88_F | CCGG TGGTAGCTAGTTGTTTCAGAA CTCGAG TTCTGAAACAACTAGCTACCA TTTTTG | shRNA |
| shIFT88_R | AATTCAAAAA TGGTAGCTAGTTGTTTCAGAA CTCGAG TTCTGAAACAACTAGCTACCA | shRNA |
| Kif24_TS_F | GCTGCACCTAAGGTCTCTGG | Mutagenesis |
| Kif24_TS_R | AGCAAAAGGAATGTTGGGTG | Mutagenesis |
| H2BC11_F | AATCTCGAGGCCACCATGCCAGAGCCAGCGAAGTC | Subcloning |
| H2BC11_R | AATCTCGAGCTTAGCGCTGGTGTACTTGG | Subcloning |
| Kif24-primer1_F | GGAGGTACGTCGTGGAGAAA | qPCR |
| Kif24-primer1_R | CGCCTCACCAAAGACTTCAT | qPCR |
| Kif24-primer2_F | CTTGGCTGGCAGTGAAAGAG | qPCR |
| Kif24-primer2_R | TGTGTTCCTGATCCAGTGCT | qPCR |
| hGAPDH_F | GGCTGAGAACGGGAAGCTTG | qPCR |
| hGAPDH_R | ACTCCACGACGTACTCAGCG | qPCR |

**Table S1**
